# Structural basis of the Integrator complex assembly and association with transcription factors

**DOI:** 10.1101/2024.01.30.577955

**Authors:** Michal Razew, Angelique Fraudeau, Moritz M. Pfleiderer, Wojciech P. Galej

## Abstract

Integrator is a multi-subunit protein complex responsible for premature transcription termination of coding and non-coding RNAs in Metazoans. This is achieved via Integrator’s two enzymatic activities, RNA endonuclease and protein phosphatase, acting on the promoter-proximally paused RNA Polymerase II (RNAPII). Yet, it remains unclear how Integrator assembly and recruitment are regulated and what are the functions of many of its core subunits. Here we report two cryo-EM reconstructions of large Integrator sub-complexes: INTS10/13/14/15 (Arm module) and INTS5/8/10/15, which allowed integrative modelling of the fully-assembled Integrator bound to the RNAPII paused elongating complex (PEC). INTS13/14 are positioned near the DNA upstream from the transcription pause site, suggesting a potential role in the chromatin context. An *in silico* protein interaction screen of over 1500 transcription factors (TFs), identified Zinc Finger Protein 655 (ZNF655) as a direct interacting partner of INTS13 that associates with a fully assembled, 17-subunit Integrator complex. We propose a model wherein the Arm module acts as a platform for the recruitment of TFs that could modulate the stability of the Integrator’s association at specific loci and modulate transcription attenuation of the target genes.

## Introduction

In metazoans, one of the critical layers of transcription regulation involves promoter-proximal pausing of the RNA polymerase II (RNAPII), which is controlled by the association of RNAPII with negative regulators of transcription DSIF and NELF^1^. Recruitment of the P-TEFb complex and its kinase activity triggers the release of the paused polymerase into the gene body and allows for productive transcription elongation^2^. However, not all stalled RNAPII are destined to produce mature transcripts, and premature transcription termination can be initiated by the recruitment of the multi-subunit Integrator complex to the paused polymerase^3–10^. Integrator complex was discovered initially as the 3’-end processing endonuclease of the non-polyadenylated snRNA transcripts^3^. More recent studies pointed to its emerging role in several aspects of the transcription of protein-coding genes and other non-coding RNAs^11–18^. Integrator’s association with the protein phosphatase PP2A-C counteracts CDK9 kinase of the P-TEFb complex and provides compelling evidence that a fine balance between these two activities plays a key role in determining the outcome of gene expression^19–22^. Consequently, perturbations of the Integrator function are known to result in severe genetic disorders^23–25^.

Integrator complex assembles from fifteen canonical subunits (INTS1-15), including recently uncovered INTS15^26–30^ and together with PP2A-C heterodimer forms a 1.5 MDa Integrator-PP2A complex that harbours endonuclease (INTS11) and phosphatase (PP2A-C) enzymatic activities^31–34^. Numerous other factors have been reported to interact with the Integrator, most notably transcription regulators (e.g. PAF1C), transcription factors (TFs) (eg. ZNF592, ZNF687, ZFP609) and chromatin reading/modifying enzymes (e.g. ZMYND8)^35–37^.

Biochemical and structural studies revealed several stable sub-complexes of the Integrator, including the *Cleavage Module* (INTS4/9/11) containing β-CASP/MBL catalytic endonuclease INTS11^38–40^, as well as the INTS10/13/14 module^41,42^ and its extended variant INTS10/13/14/15^27,29^ referred to as the *Arm module* hereafter. Cryo-EM reconstructions of the higher-order Integrator-PP2A complexes revealed its overall architecture and intimate interactions between different modules as well as the structural basis of its recruitment to the paused elongating RNAPII complex (PEC)^43^ and the mechanism of INTS11 endonuclease activation^20,44,45^.

Despite the tremendous progress that has been made in structural studies of the Integrator, many of its subunits, including the entire Arm (INTS10/13/14/15) module, have never been resolved in the cryo-EM studies and their role in the context-dependent function of the integrator is not well understood. It is also not clear how TFs or chromatin factors associate with the Integrator and what their role is in Integrator recruitment to target genes.

Here we report cryo-EM structures of two Integrator sub-complexes and a structural model of a fully assembled 17-subunit Integrator complex bound to PEC. This, together with AlphaFold2-based high-throughput *in silico* screening of protein-protein interactions, revealed the structural basis for the Integrator interaction with a zinc finger transcription factor ZNF655. Our case study may serve as a more general model for understanding the mechanism of Integrator association with a broader spectrum of adaptor proteins.

## Results

### cryo-EM structure of the Integrator Arm module

Integrator Arm module containing subunits INTS10/13/14/15 has been previously characterised biochemically^27,29^, but experimental data supporting its overall architecture remained missing. We expressed the 270 kDa INTS10/13/14/15 quaternary complex recombinantly in insect cells (Figure 1 and Figure S1A-B) and determined its structure using single-particle cryoEM at 3.3 Å resolution (Figure 1A and B; Figure S2; Table 1). The overall architecture of the Arm module is elongated, hook-shaped with INTS15 and INTS13/14 separated from each other by nearly 200 Å and connected exclusively by an extended and mostly alpha-helical INTS10 (Figure 1). INTS15 interacts with INTS10 in a head-to-tail arrangement, and a weaker cryo-EM density in this region suggests potential conformational dynamics (Figure 1). INTS14^VWA^ domain coordinates a metal ion at the interface with INTS10, which is also bound by E633 of INTS10 (Figure 1E). This interaction resembles previously reported Mg^2+^-mediated ICAM-3 and integrin α_L_β_2_ interface^46^. Binary interactions formed by the components of the Arm module are in good agreement with the previously proposed integrative model^29^ and the crystal structure of INTS13/14^42^ (Figure S3).

**Figure 1.**
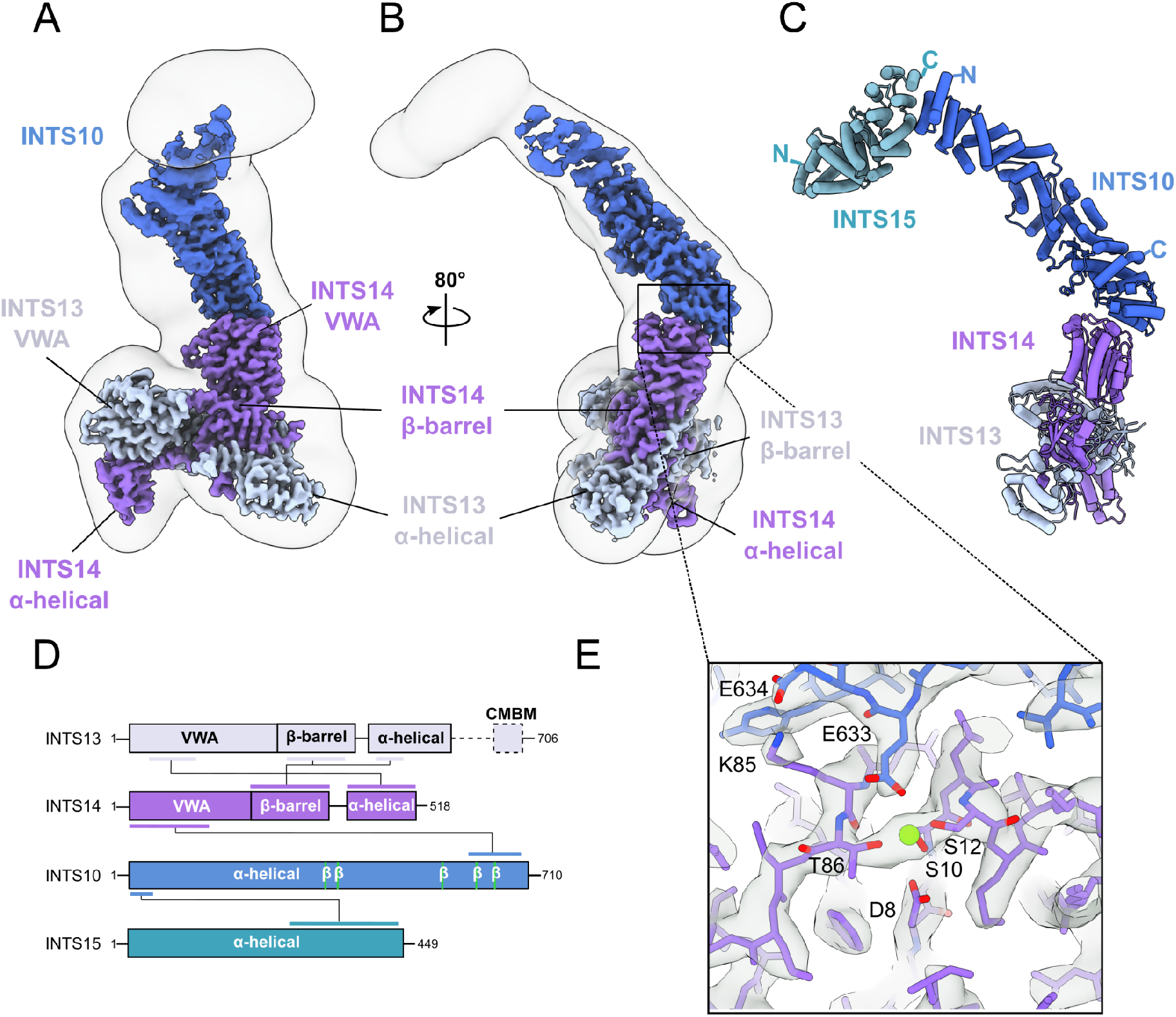
CryoEM structure of the INTS10/13/14/15 complex (Arm module). **A)** High-resolution cryoEM map coloured by the subunit identity fitted in a low-contour level map showing less well-ordered, peripheral regions. **B)** orthogonal view of the complex. **C)** Atomic model of the Integrator Arm module. **D)** Schematic of the domain composition of each subunit of the Integrator Arm module. Coloured lines indicate interacting regions. The dashed line in INTS13 indicates regions that are missing in our density, including the Cleavage Module Binding Motif (CMBM). **E)** Zoom in on the INTS10-INTS14 interface showing amino acid residues fitting into the cryoEM density. The metal ion within the Metal-Ion Dependent Adhesion Site (MIDAS) of the INTS14 VWA domain is highlighted as a green sphere.

### Interaction of the Arm module with the core of the Integrator complex

While we obtained insights into the structure of the isolated Integrator Arm module, it remained unclear how it interacts with other Integrator components within the fully assembled complex. Two contacts between the Arm module and the Integrator’s core have been previously reported: Cleavage Module Binding Motif (CMBM) located at the C-terminus of INTS13 interaction with INTS4/9/11^42,47^ and INTS5-INTS15 interaction predicted using AlphaFold2 interaction screen^29^. The molecular basis for both of these interactions remains unknown.

To directly address this knowledge gap, we reconstituted a quaternary complex of INTS5/8/10/15 and determined its structure by cryo-EM at the overall 3.2 Å resolution (Figure 2). High anisotropy of the reconstruction does not allow for accurate side-chain modelling, therefore the map was only used after low-pass filtering for fitting in secondary structure elements (Figure S4). The best-defined region of the cryo-EM map is centred around INTS5/8 (Figure 2A), which remains unchanged when compared to previously reported structures of the Integrator-PP2A^20^ and Integrator-PEC complexes^44,45^. The map quality gradually decreases towards the region encompassing INTS15 and INTS10 (Figure 2A). Although the limited resolution did not allow for the *de novo* building of the atomic models of INTS10 and INTS15, the structures of both proteins could be unambiguously fitted into the map based on the Arm module reconstruction (Figure 2B). INTS5/8/10/15 complex has an elongated architecture spanning nearly 300 Å in the longest dimension. INTS15 plays a key role in bridging INTS5/8 and INTS10, which otherwise make no contact with one another.

**Figure 2.**
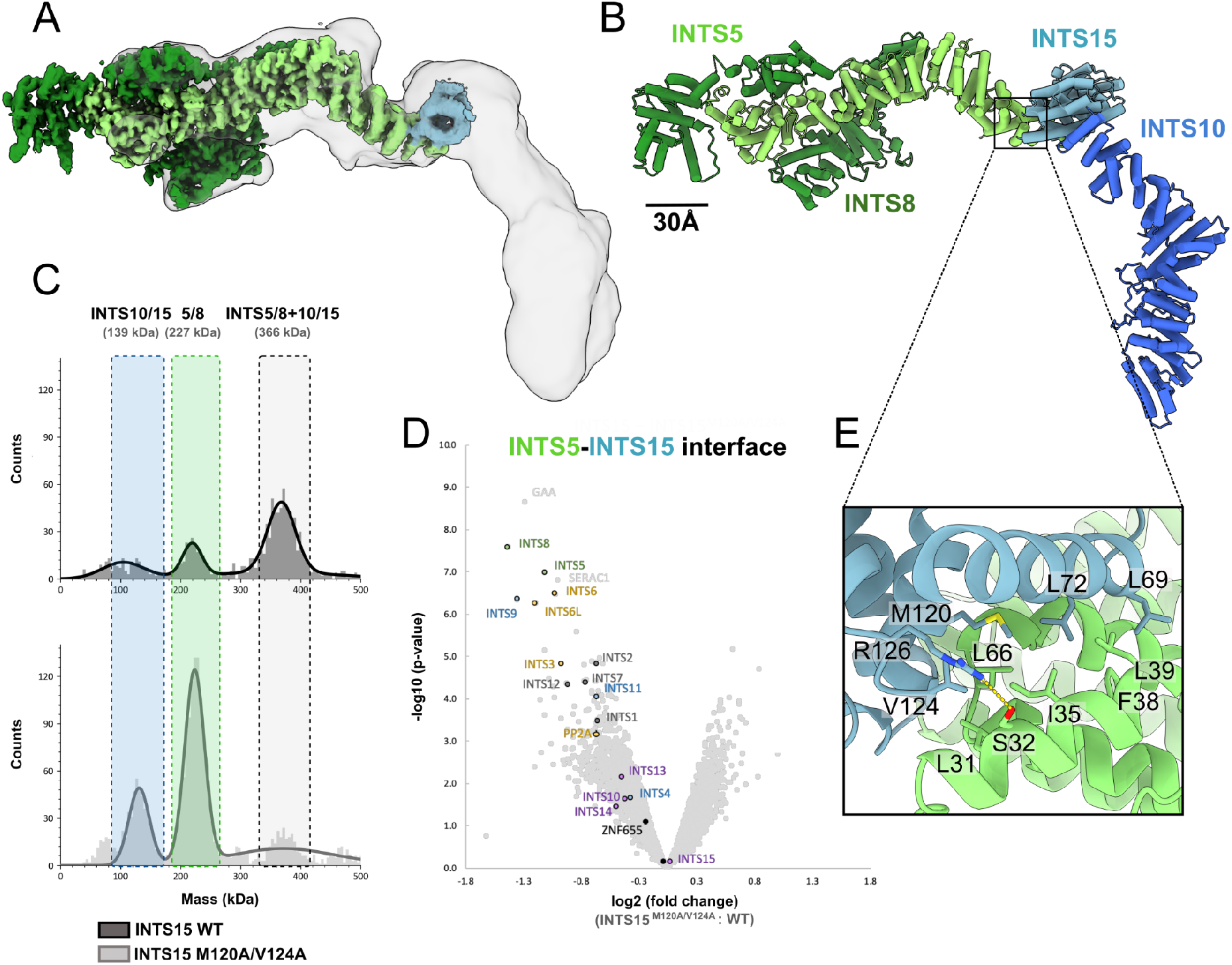
CryoEM structure of the INTS5/8/10/15 complex. **A)** High-resolution cryoEM map coloured by the subunit identity fitted in a low-contour level map showing less well-ordered peripheral regions. **B)** Integrative atomic model of the INTS5/8/10/15 complex based on experimental cryoEM density and AF2 model **C)** Mass photometry analysis of the INTS5/8/10/15 complex formation from recombinant proteins containing either a wild-type INTS15 or its M102A/V124A variant predicted to abolish complex formation **D)** Differential quantitative mass spectrometry analysis of the proteins co-purifying with 3xHA-INTS15(WT or M102A/V124A) transiently expressed in HEK293T cells and purified on Anti-HA-agarose resin. **E)** Zoom-in at the interface of INTS5-INTS15 modelled with AlphaFold2 and fitted into low-resolution cryoEM reconstruction.

To validate the INTS5-INTS15 interface observed in our cryo-EM reconstruction (Figure 2E), we designed point mutation variants of INTS15 (INTS15^M120A/V124A^, INTS15^L69A/L72A^ and INTS15^C157P/C158P^) expected to abolish this interaction and analysed their effects *in vitro* and *in vivo*. First, we transiently expressed INTS15 and its variants in HEK293T cells, purified them using hemagglutinin (HA)-agarose resin and subjected them to TMT-plex differential quantitative proteomics (Figure 2D). When compared to the INTS15^WT^, both the M120A/V124A and L69A/L72A variants have lost their ability to co-purify numerous Integrator subunits (Figure 2D, Figure S6A). These include INTS5, a direct interacting partner of INTS15, which, together with INTS8, was amongst the highest-ranking hits. Similarly, enrichments of the phosphatase (PP2A-INTS6) and backbone modules (INTS1/2/7/12) were decreased in the INTS15^M120A/V124A^ and INTS15^L69A/L72A^ mutants, which is to be expected as both of these modules are known to interact with INTS5/8^20^. Notably, no significant changes to the abundance of the INTS15 and other components of the Arm module were observed, as those interactions were not targeted by the mutations discussed.

Next, we purified recombinant INTS5/8 and INTS10/15 (WT and M120A/V124A) and analysed their association *in vitro* using mass photometry^48^. When INTS15^WT^ was used, we observed an efficient formation of a large complex with the measured molecular weight corresponding very well to the theoretical mass of INTS5/8/10/15 (367 kDa vs 366 kDa), while the INTS15^M120A/V124A^ mutant severely hampered complex formation (Figure 2C and Supplementary Table 1).

Taken together, these data establish the molecular basis of the INTS5-INTS15 interaction and provide new insights into the mode of the Arm module recruitment to the core of the Integrator.

### Model of the fully assembled Integrator complex bound to RNAPII

Integrator bound to paused elongating RNAPII forms a compact structure with multiple contacts between RNAPII and Integrator subunits, including INTS1, INTS6, INTS7 and INTS11^44,45^. Negative regulators of transcription DSIF, NELF play critical roles in the Integrator recruitment and activation of its endonuclease INTS11. The structure of the 4-subunit Arm module has never been visualised in the context of the fully assembled Integrator. To address this issue, we first superimposed the two cryo-EM structures presented here via overlapping subunits, INTS10/15, to create a model of the INTS5/8/10/13/14/15 complex (Figure 3A). This structure was then oriented with respect to the Integrator-PEC complex by superposition via INTS5, a subunit well resolved in current and previous studies^44,45^. Re-analysis of the previously deposited cryo-EM maps of the Integrator-PP2A complexes (EMD-33741 and EMD-30473)^20,45^ (Figure 3B, Figure S7) shows low-resolution unassigned density, which is in good agreement with the location of the Arm module obtained by the superposition (Figure 3C). This density has not been interpreted previously, most likely due to the limited resolution of this region of the map and missing high-resolution information for the Arm module presented here. The positioning of the Arm module is also compatible with the previously reported interaction between INTS13^CMBM^ and the Cleavage module^42,47^. Interestingly, the EMD-33741 map contains additional density near the CTD1 domain of the INTS9/11 heterodimer, which is in excellent agreement with the INTS13^CMBM^ interaction site predicted by AF2 (Figure S8A-D).

**Figure 3.**
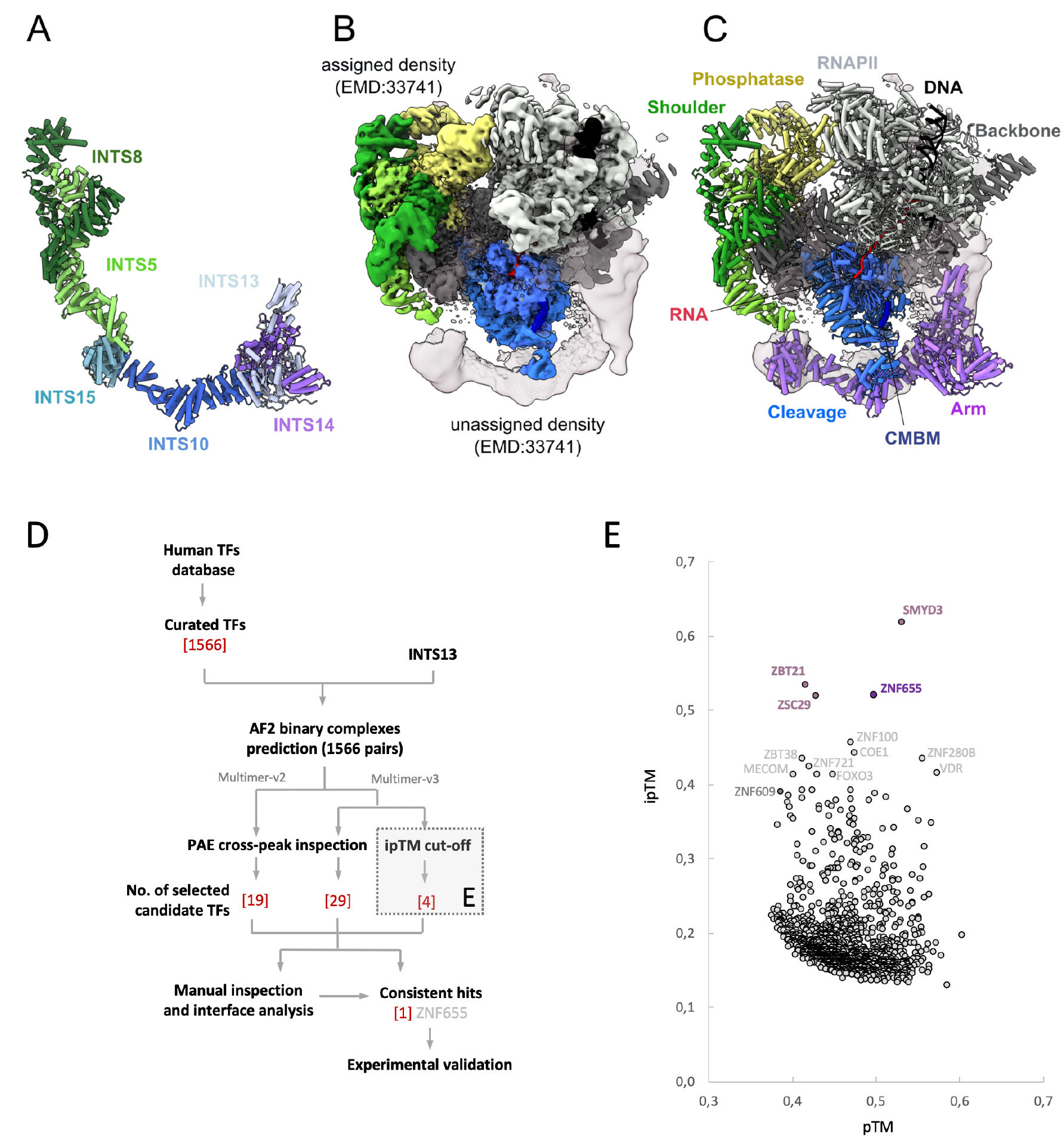
Model of the fully assembled Integrator complex bound to paused elongating RNAPII. **A)** A composite model of the Integrator Shoulder-Arm module (INTS5/8/10/13/14/15) based on two experimental cryoEM structures aligned using shared subunits (INTS10/15). **B)** CryoEM map of the Integrator-RNAPII structure (EMD: 33741) contoured at 3.2σ, coloured by regions assigned to RNAPII and Integrator modules. The unassigned density was contoured at 0.25σ. **C)** The atomic model of the fully assembled Integrator-RNAPII with the same orientation and colouring as in B. Cleavage Module Binding Motif of INTS13 (CMBM) was identified based on the AlphaFold2 prediction. The Integrator Shoulder-Arm module from panel A was superimposed on the Integrator-PEC structure via INTS5. The position of the Arm module (violet) fits very well to the unassigned density. **D)** A workflow of the *in silico* protein-protein interaction screen used in this study. **E)** Predicted template modelling score (pTM) and interface predicted template modelling score (ipTM) for INTS13-TFs pairs analysed with AlphaFold multimer v3. TFs with the ipTM values above 0.4 were labelled explicitly and the top 4 hits were coloured in pink/magenta.

Our new data, together with previous cryo-EM data, allow unambiguous modelling of the Arm module in association with the Integrator-PEC complex. One striking feature of this model is the location of the Arm module in the proximity of the DNA upstream from the transcription pause site, which would in principle, allow their direct or indirect interaction (Figure S8E-F). We hypothesised that based on its location, INTS13/14 might act as a platform to recruit additional proteins, such as Transcription Factors (TFs), that might be involved in mediating Integrator’s interactions with DNA. Therefore, we set off to investigate the structural basis of the Integrator association with selected TFs.

### Integrator association with transcription factors

Previous studies have shown that Integrator interacts with various transcription factors (TFs), including the so-called Z3 complex composed of ZNF592, ZNF687 and ZMYND8^35,49^; ZFP609^35,36^ and ZNF655^29^. However, the structural basis of these interactions remains elusive.

In order to identify a more comprehensive set of Integrator-binding TFs, we used AlphaFold2 (AF2)^50,51^ as a tool to screen protein-protein interactions. We used a self-curated version of the database of all human transcription factors^52^ and performed high-throughput structure prediction of over 1500 TFs paired with INTS13 (Figure 3D). Based on intermolecular PAE cross-peaks from the two independent runs using different version of the AF2 model and predicted interface template modelling scores (ipTM), we identified a small subset of TFs predicted with high confidence to bind to INTS13 (Figure 3E). One of these hits, ZNF655, is a 491AA protein belonging to the Kruppel-like family of transcription factors containing 6 C2H2 type zinc finger domains at the C-terminus^53^ (Figure 4A), known to be upregulated in human pancreatic cancers^54^. We showed previously that ZNF655 indeed co-purifies with INTS13/14^29^, therefore, we focussed on the characterisation of the ZNF655-INTS13 interaction in more detail.

**Figure 4.**
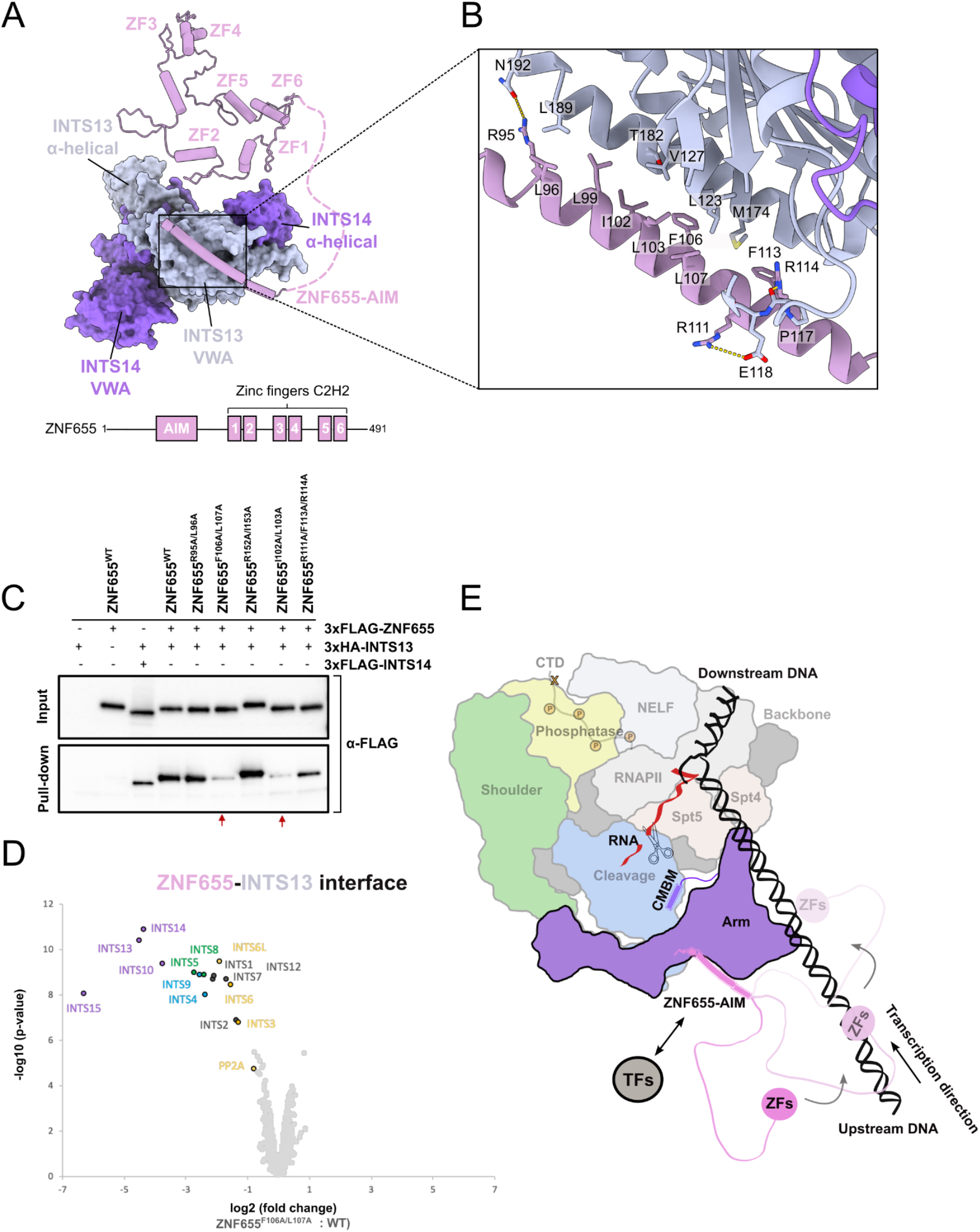
Transcription factor ZNF655 interacts with the Integrator complex via its Arm module interacting motif. **A)** Structural model of ZNF655 bound to INTS13/14 of the Integrator Arm module based on the AlphaFold2 prediction and a schematic of ZNF655 domain organisation. AIM -Arm Interacting Motif; ZF1-6 - Zinc Fingers 1-6. **B)** Zoom in on the AlphaFold2 predicted ZNF655^AIM^-INTS13^VWA^ interface, coloured as in A. **C)** ZNF655 mutagenesis and INTS13 pull-down experiment validating the interface described in B. ZNF655 variants were used as prey and INTS13 as bait. INTS14 was used as a positive control, and ZNF655^R152A/R153A^ as a negative control. **D)** Differential quantitative proteomics experiment of the 3xFLAG-ZNF655^F106A/L107A^ variant and 3xFLAG-ZNF655^WT^ transiently expressed in HEK293T cells and purified on anti-FLAG affinity resin. **E)** Schematic model of ZNF655 bound to Integrator-PEC complex. ZNF655 zinc fingers (ZFs) are connected to the Arm Interacting Motif (AIM) via a long, disordered linker which enables it to probe the surroundings for DNA binding sites (see also Figure S10).

### ZNF655 interaction with the fully assembled Integrator complex

First, we reversed the AF2 screening procedure to look for binary interactions between ZNF655 and all known Integrator subunits. Based on intermolecular PAE cross-peaks, we identified INTS13 as the sole hit and the most likely binding partner of ZNF655, consistent with the result of the initial screen (Figure S10A-C). The predicted interface is formed between the N-terminal helix of ZNF655 and a hydrophobic cleft in the VWA domain of INTS13 (Figure 4A-B). To validate this interaction, we designed a series of point mutations in ZNF655 at the INTS13 interface and performed a pull-down experiment from HEK293T cells transiently co-expressing 3xFLAG-ZNF655 (WT or variants) and 3xHA-INTS13. Proteins associated with the HA-agarose beads were analysed by western blot against the FLAG-tag to probe for the presence of ZNF655 (Figure 4C). While INTS13 could efficiently enrich ZNF655^WT^, the interaction was nearly completely lost in the case of ZNF655 double-mutants of interface residues: F106A/I107A and I102A/I103A (Figure 4C). ZNF655^R152A/I153A^ was used as a negative control (residues not located at the interface).

Although we established that ZNF655 interacts specifically with INTS13, a broader functional context of this association remains unclear (i.e. if this binding concerns only the Arm module or also the fully assembled Integrator). To address this, we compared the proteome of ZNF655^WT^ with ZNF655^F106A/L107A^ mutant deficient in INTS13 interaction (Figure 4D). Disruption of the ZNF655-INTS13 interface leads to the loss of not only INTS13 and the Arm module subunits, but essentially the entire Integrator complex (Figure 4D). This provides evidence that ZNF655 associates stably with the fully assembled Integrator complex.

ZNF655 interaction with INTS13 established here implies that its zinc fingers would be loosely tethered to Integrator via an 82 AA disordered linker, and could potentially probe their surroundings within a radius of approximately 260 Å (equivalent of +/-80bp from the TSS) (Figure 4E and Figure S10D). That would allow them to reach the DNA upstream or downstream from the transcription pause site, as well as nascent RNA. Therefore ZNF655 zinc fingers could be, in principle, utilised to modulate the stability of Integrator’s association with the paused RNAPII.

## Discussion

In this report, we present two cryo-EM structures of large Integrator sub-complexes, INTS5/8/10/15 and INTS10/13/14/15 (Arm module), which, together with the previously deposited cryo-EM data, allowed us to create the most complete model of the Integrator complex assembled in the context of the promoter-proximally paused RNAPII. Our model revealed the location of the previously not resolved subunits INTS10/13/14/15 with respect to the core Integrator and RNAPII. INTS15 was identified as the pivotal factor linking INTS13/14 heterodimer with the core of the Integrator via the extended alpha-helical structure of INTS10. INTS13/14 are positioned in the path of the upstream DNA extrapolated from the RNAPII structure^44,45^. INTS13/14 has been shown to have weak and not specific affinity for the nucleic acids^42^, but the functional implications of this observation remained unclear. Our model suggests that these subunits could be indeed involved in direct or indirect interaction with DNA, and their positioning by the fully assembled Integrator could explain weak DNA binding properties when tested in isolation.

We hypothesised that nucleic acid binding properties of INTS13/14 could be supported by additional proteins, including transcription factors. We used AlphaFold2 (AF2)^50^ to exhaustively screen *in silico* all possible interactions between Integrator subunit INTS13 and all annotated human TFs^52^. We previously showed on a much smaller scale that such a screening approach can uncover new and unexpected protein-protein interactions^29^. Here, our screen identified a small subset of TFs predicted to form high-confidence interfaces with INTS13. As a proof of concept, we performed a detailed characterisation of one of them, ZNF655, whose interaction with INTS13 was confirmed experimentally. This successful example shows the potential of a more general application of AF2-based interaction screening approaches that may aid future biochemical and structural studies.

Our data provide the first example of a structural basis for the association of Integrator with a zinc finger transcription factor^55^. The binding mode of ZNF655 to INTS13 is compatible with the fully assembled Integrator bound to PEC, and we show experimentally that ZNF655 co-purifies with virtually all Integrator subunits. At present, there is no clear biochemical evidence supporting ZNF655’s association with DNA, chromatin or RNA, as shown for some TFs containing RNA binding motifs^56^. However, ZNF655 was found in a screen for RNA-binding regions within the nuclear proteome of embryonic stem cells^57^ and several studies reported its links to cell proliferation in various types of cancer^54,58,59^, supporting its potential role as a transcription regulator. Future studies of ZNF655 DNA/RNA binding abilities and its chromatin localization are necessary to better understand its regulatory functions. Given several other TFs are known to associate with the Integrator, the example of ZNF655 described here may represent a more general paradigm for Integrator interaction with other adaptor proteins. It remains to be seen how other TFs and/or chromatin-binding proteins are associated with the Integrator and if their combinatorial or mutually exclusive association can play a functional role in the Integrator targeting.

Integrator association with TFs points to the possibility of its targeting to specific loci through intrinsic sequence specificities of TFs^60^. Early work on snRNA processing suggested possible specific Integrator recruitment to the snRNA gene loci via its interaction with the SNAPc and so-called 3’-box located in pre-snRNA transcripts^61^. However, more recent findings show that Integrator acts on diverse classes of RNAs, and no clear consensus sequences neither in the DNA around Integrator binding sites nor in the RNA transcripts near its cleavage sites, could be identified^34^. Therefore a universal mechanism wherein the Integrator would be directed/instructed in a sequence-specific manner seems unlikely.

Here we envisage that the Integrator’s interaction with TFs could be utilised for fine-tuning and modulation of the Integrator binding stability and/or dwell time at the transcription pause sites rather than as a means of specific recruitment. Altering the stability of Integrator association with RNAPII at particular loci would shift the force balance in the PP2A-CDK9 axis and consequently alter the output of gene expression. Such context-specific regulation by TFs would be compatible with numerous reports on the Integrator-mediated rapid response to environmental stress and signal induction^18,62,63^.

## Acknowledgements

We would like to acknowledge Mandy Rettel, Per Haberkant, and Frank Stein from the EMBL Proteomic Core Facility for their support with the mass spectrometry data acquisition and analysis; Romain Linares for the assistance with the cryo-EM data collection; Sarah Schneider and Joe Bartho for ensuring smooth running of the EMBL cryo-EM facilities; Magdalena Stoykova, Yonca Ozturk and Tara Best for the technical support. Martin Pelosse from the EMBL Grenoble EEF platform for assistance with cell culture. A. Peuch (IBS) and the EMBL Grenoble IT team for the support with high-performance computing. Sebastian Eustermann for critical comments on the manuscript. This work used the platforms of the Grenoble Instruct-ERIC centre (ISBG; UAR 3518 CNRS-CEA-UGA-EMBL) within the Grenoble Partnership for Structural Biology (PSB), supported by FRISBI (ANR-10-INBS-0005-02) and GRAL, financed within the University Grenoble Alpes graduate school (Ecoles Universitaires de Recherche) CBH-EUR-GS (ANR-17-EURE-0003). We thank Caroline Mas for access to the biophysics platform. M.R. was supported by a fellowship from the EMBL Interdisciplinary Postdoc (EIPOD) programme under Marie Sklodowska-Curie Actions COFUND (grant agreement no. 847543).

## Author contributions

M.R. expressed and purified recombinant proteins, performed mass photometry experiments, prepared cryoEM grids, collected and analysed cryo-EM data and built corresponding models. A.F. performed pull-down, western blotting and sample preparation for quantitative proteomics. M.M.P. Generated insect cell expression constructs and optimised recombinant expression and purification of the integrator subunits. W.P.G. designed and performed the AF2 screen and supervised the project. All authors analysed and interpreted data. M.R. and W.P.G. wrote the manuscript with input from all authors.

## References

1. Adelman, K., and Lis, J.T. (2012). Promoter-proximal pausing of RNA polymerase II: emerging roles in metazoans. Nat. Rev. Genet. 13, 720–731.

2. Peterlin, B.M., and Price, D.H. (2006). Controlling the elongation phase of transcription with P-TEFb. Mol. Cell 23, 297–305.

3. Baillat, D., Hakimi, M.-A., Näär, A.M., Shilatifard, A., Cooch, N., and Shiekhattar, R. (2005). Integrator, a multiprotein mediator of small nuclear RNA processing, associates with the C-terminal repeat of RNA polymerase II. Cell 123, 265–276.

4. Gardini, A., Baillat, D., Cesaroni, M., Hu, D., Marinis, J.M., Wagner, E.J., Lazar, M.A., Shilatifard, A., and Shiekhattar, R. (2014). Integrator regulates transcriptional initiation and pause release following activation. Mol. Cell 56, 128–139.

5. Stadelmayer, B., Micas, G., Gamot, A., Martin, P., Malirat, N., Koval, S., Raffel, R., Sobhian, B., Severac, D., Rialle, S., et al. (2014). Integrator complex regulates NELF-mediated RNA polymerase II pause/release and processivity at coding genes. Nat. Commun. 5, 5531.

6. Yamamoto, J., Hagiwara, Y., Chiba, K., Isobe, T., Narita, T., Handa, H., and Yamaguchi, Y. (2014). DSIF and NELF interact with Integrator to specify the correct post-transcriptional fate of snRNA genes. Nat. Commun. 5, 4263.

7. Elrod, N.D., Henriques, T., Huang, K.-L., Tatomer, D.C., Wilusz, J.E., Wagner, E.J., and Adelman, K. (2019). The Integrator Complex Attenuates Promoter-Proximal Transcription at Protein-Coding Genes. Mol. Cell 76, 738–752.e7.

8. Tatomer, D.C., Elrod, N.D., Liang, D., Xiao, M.-S., Jiang, J.Z., Jonathan, M., Huang, K.-L., Wagner, E.J., Cherry, S., and Wilusz, J.E. (2019). The Integrator complex cleaves nascent mRNAs to attenuate transcription. Genes Dev. 33, 1525–1538.

9. Lykke-Andersen, S., Žumer, K., Molska, E.Š., Rouvière, J.O., Wu, G., Demel, C., Schwalb, B., Schmid, M., Cramer, P., and Jensen, T.H. (2021). Integrator is a genome-wide attenuator of non-productive transcription. Mol. Cell 81, 514–529.e6.

10. Stein, C.B., Field, A.R., Mimoso, C.A., Zhao, C., Huang, K.-L., Wagner, E.J., and Adelman, K. (2022). Integrator endonuclease drives promoter-proximal termination at all RNA polymerase II-transcribed loci. Mol. Cell 82, 4232–4245.e11.

11. Cazalla, D., Xie, M., and Steitz, J.A. (2011). A primate herpesvirus uses the integrator complex to generate viral microRNAs. Mol. Cell 43, 982–992.

12. Lai, F., Gardini, A., Zhang, A., and Shiekhattar, R. (2015). Integrator mediates the biogenesis of enhancer RNAs. Nature 525, 399–403.

13. Beckedorff, F., Blumenthal, E., daSilva, L.F., Aoi, Y., Cingaram, P.R., Yue, J., Zhang, A., Dokaneheifard, S., Valencia, M.G., Gaidosh, G., et al. (2020). The Human Integrator Complex Facilitates Transcriptional Elongation by Endonucleolytic Cleavage of Nascent Transcripts. Cell Rep. 32, 107917.

14. Dasilva, L.F., Blumenthal, E., Beckedorff, F., Cingaram, P.R., Gomes Dos Santos, H., Edupuganti, R.R., Zhang, A., Dokaneheifard, S., Aoi, Y., Yue, J., et al. (2021). Integrator enforces the fidelity of transcriptional termination at protein-coding genes. Sci Adv 7, eabe3393.

15. Kirstein, N., Dokaneheifard, S., Cingaram, P.R., Valencia, M.G., Beckedorff, F., Gomes Dos Santos, H., Blumenthal, E., Tayari, M.M., Gaidosh, G.S., and Shiekhattar, R. (2023). The Integrator complex regulates microRNA abundance through RISC loading. Sci Adv 9, eadf0597.

16. Beltran, T., Pahita, E., Ghosh, S., Lenhard, B., and Sarkies, P. (2021). Integrator is recruited to promoter-proximally paused RNA Pol II to generate Caenorhabditis elegans piRNA precursors. EMBO J. 40, e105564.

17. Rubtsova, M.P., Vasilkova, D.P., Moshareva, M.A., Malyavko, A.N., Meerson, M.B., Zatsepin, T.S., Naraykina, Y.V., Beletsky, A.V., Ravin, N.V., and Dontsova, O.A. (2019). Integrator is a key component of human telomerase RNA biogenesis. Sci. Rep. 9, 1701.

18. Rosa-Mercado, N.A., Zimmer, J.T., Apostolidi, M., Rinehart, J., Simon, M.D., and Steitz, J.A. (2021). Hyperosmotic stress alters the RNA polymerase II interactome and induces readthrough transcription despite widespread transcriptional repression. Mol. Cell 81, 502–513.e4.

19. Huang, K.-L., Jee, D., Stein, C.B., Elrod, N.D., Henriques, T., Mascibroda, L.G., Baillat, D., Russell, W.K., Adelman, K., and Wagner, E.J. (2020). Integrator Recruits Protein Phosphatase 2A to Prevent Pause Release and Facilitate Transcription Termination. Mol. Cell 80, 345–358.e9.

20. Zheng, H., Qi, Y., Hu, S., Cao, X., Xu, C., Yin, Z., Chen, X., Li, Y., Liu, W., Li, J., et al. (2020). Identification of Integrator-PP2A complex (INTAC), an RNA polymerase II phosphatase. Science 370. 10.1126/science.abb5872.

21. Vervoort, S.J., Welsh, S.A., Devlin, J.R., Barbieri, E., Knight, D.A., Offley, S., Bjelosevic, S., Costacurta, M., Todorovski, I., Kearney, C.J., et al. (2021). The PP2A-Integrator-CDK9 axis fine-tunes transcription and can be targeted therapeutically in cancer. Cell 184, 3143–3162.e32.

22. Hu, S., Peng, L., Song, A., Ji, Y.-X., Cheng, J., Wang, M., and Chen, F.X. (2023). INTAC endonuclease and phosphatase modules differentially regulate transcription by RNA polymerase II. Mol. Cell 83, 1588–1604.e5.

23. Oegema, R., Baillat, D., Schot, R., van Unen, L.M., Brooks, A., Kia, S.K., Hoogeboom, A.J.M., Xia, Z., Li, W., Cesaroni, M., et al. (2017). Human mutations in integrator complex subunits link transcriptome integrity to brain development. PLoS Genet. 13, e1006809.

24. Tilley, F.C., Arrondel, C., Chhuon, C., Boisson, M., Cagnard, N., Parisot, M., Menara, G., Lefort, N., Guerrera, I.C., Bole-Feysot, C., et al. (2021). Disruption of pathways regulated by Integrator complex in Galloway–Mowat syndrome due to WDR73 mutations. Sci. Rep. 11, 1–13.

25. Tepe, B., Macke, E.L., Niceta, M., Weisz Hubshman, M., Kanca, O., Schultz-Rogers, L., Zarate, Y.A., Schaefer, G.B., Granadillo De Luque, J.L., Wegner, D.J., et al. (2023). Bi-allelic variants in INTS11 are associated with a complex neurological disorder. Am. J. Hum. Genet. 110, 774–789.

26. Drew, K., Wallingford, J.B., and Marcotte, E.M. (2021). hu.MAP 2.0: integration of over 15,000 proteomic experiments builds a global compendium of human multiprotein assemblies. Mol. Syst. Biol. 17, e10016.

27. Replogle, J.M., Saunders, R.A., Pogson, A.N., Hussmann, J.A., Lenail, A., Guna, A., Mascibroda, L., Wagner, E.J., Adelman, K., Lithwick-Yanai, G., et al. (2022). Mapping information-rich genotype-phenotype landscapes with genome-scale Perturb-seq. Cell 185, 2559–2575.e28.

28. Pan, J., Kwon, J.J., Talamas, J.A., Borah, A.A., Vazquez, F., Boehm, J.S., Tsherniak, A., Zitnik, M., McFarland, J.M., and Hahn, W.C. (2022). Sparse dictionary learning recovers pleiotropy from human cell fitness screens. Cell Syst 13, 286–303.e10.

29. Offley, S.R., Pfleiderer, M.M., Zucco, A., Fraudeau, A., Welsh, S.A., Razew, M., Galej, W.P., and Gardini, A. (2023). A combinatorial approach to uncover an additional Integrator subunit. Cell Rep. 42, 112244.

30. Azuma, N., Yokoi, T., Tanaka, T., Matsuzaka, E., Saida, Y., Nishina, S., Terao, M., Takada, S., Fukami, M., Okamura, K., et al. (2023). Integrator complex subunit 15 controls mRNA splicing and is critical for eye development. Hum. Mol. Genet. 32, 2032–2045.

31. Pfleiderer, M.M., and Galej, W.P. (2021). Emerging insights into the function and structure of the Integrator complex. Transcription 12, 251–265.

32. Sabath, K., and Jonas, S. (2022). Take a break: Transcription regulation and RNA processing by the Integrator complex. Curr. Opin. Struct. Biol. 77, 102443.

33. Welsh, S.A., and Gardini, A. (2023). Genomic regulation of transcription and RNA processing by the multitasking Integrator complex. Nat. Rev. Mol. Cell Biol. 24, 204–220.

34. Wagner, E.J., Tong, L., and Adelman, K. (2023). Integrator is a global promoter-proximal termination complex. Mol. Cell 83, 416–427.

35. Malovannaya, A., Lanz, R.B., Jung, S.Y., Bulynko, Y., Le, N.T., Chan, D.W., Ding, C., Shi, Y., Yucer, N., Krenciute, G., et al. (2011). Analysis of the human endogenous coregulator complexome. Cell 145, 787–799.

36. van den Berg, D.L.C., Azzarelli, R., Oishi, K., Martynoga, B., Urbán, N., Dekkers, D.H.W., Demmers, J.A., and Guillemot, F. (2017). Nipbl Interacts with Zfp609 and the Integrator Complex to Regulate Cortical Neuron Migration. Neuron 93, 348–361.

37. Wang, Z., Song, A., Xu, H., Hu, S., Tao, B., Peng, L., Wang, J., Li, J., Yu, J., Wang, L., et al. (2022). Coordinated regulation of RNA polymerase II pausing and elongation progression by PAF1. Sci Adv 8, eabm5504.

38. Albrecht, T.R., Shevtsov, S.P., Wu, Y., Mascibroda, L.G., Peart, N.J., Huang, K.-L., Sawyer, I.A., Tong, L., Dundr, M., and Wagner, E.J. (2018). Integrator subunit 4 is a ‘Symplekin-like’ scaffold that associates with INTS9/11 to form the Integrator cleavage module. Nucleic Acids Res. 46, 4241–4255.

39. Pfleiderer, M.M., and Galej, W.P. (2021). Structure of the catalytic core of the Integrator complex. Mol. Cell 81, 1246–1259.e8.

40. Lin, M.-H., Jensen, M.K., Elrod, N.D., Huang, K.-L., Welle, K.A., Wagner, E.J., and Tong, L. (2022). Inositol hexakisphosphate is required for Integrator function. Nat. Commun. 13, 5742.

41. Barbieri, E., Trizzino, M., Welsh, S.A., Owens, T.A., Calabretta, B., Carroll, M., Sarma, K., and Gardini, A. (2018). Targeted Enhancer Activation by a Subunit of the Integrator Complex. Mol. Cell 71, 103–116.e7.

42. Sabath, K., Stäubli, M.L., Marti, S., Leitner, A., Moes, M., and Jonas, S. (2020). INTS10–INTS13–INTS14 form a functional module of Integrator that binds nucleic acids and the cleavage module. Nat. Commun. 11, 1–16.

43. Vos, S.M., Farnung, L., Urlaub, H., and Cramer, P. (2018). Structure of paused transcription complex Pol II–DSIF–NELF. Nature.

44. Fianu, I., Chen, Y., Dienemann, C., Dybkov, O., Linden, A., Urlaub, H., and Cramer, P. (2021). Structural basis of Integrator-mediated transcription regulation. Science 374, 883–887.

45. Zheng, H., Jin, Q., Wang, X., Qi, Y., Liu, W., Ren, Y., Zhao, D., Xavier Chen, F., Cheng, J., Chen, X., et al. (2023). Structural basis of INTAC-regulated transcription. Protein Cell. 10.1093/procel/pwad010.

46. Song, G., Yang, Y., Liu, J.-H., Casasnovas, J.M., Shimaoka, M., Springer, T.A., and Wang, J.-H. (2005). An atomic resolution view of ICAM recognition in a complex between the binding domains of ICAM-3 and integrin alphaLbeta2. Proc. Natl. Acad. Sci. U. S. A. 102, 3366–3371.

47. Mascibroda, L.G., Shboul, M., Elrod, N.D., Colleaux, L., Hamamy, H., Huang, K.-L., Peart, N., Singh, M.K., Lee, H., Merriman, B., et al. (2022). INTS13 variants causing a recessive developmental ciliopathy disrupt assembly of the Integrator complex. Nat. Commun. 13, 6054.

48. Young, G., Hundt, N., Cole, D., Fineberg, A., Andrecka, J., Tyler, A., Olerinyova, A., Ansari, A., Marklund, E.G., Collier, M.P., et al. (2018). Quantitative mass imaging of single biological macromolecules. Science 360, 423–427.

49. Baillat, D., Russell, W.K., and Wagner, E.J. (2016). CRISPR-Cas9 mediated genetic engineering for the purification of the endogenous integrator complex from mammalian cells. Protein Expr. Purif. 128, 101–108.

50. Jumper, J., Evans, R., Pritzel, A., Green, T., Figurnov, M., Ronneberger, O., Tunyasuvunakool, K., Bates, R., Žídek, A., Potapenko, A., et al. (2021). Highly accurate protein structure prediction with AlphaFold. Nature 596, 583–589.

51. Mirdita, M., Schütze, K., Moriwaki, Y., Heo, L., Ovchinnikov, S., and Steinegger, M. (2022). ColabFold: making protein folding accessible to all. Nat. Methods 19, 679–682.

52. Lambert, S.A., Jolma, A., Campitelli, L.F., Das, P.K., Yin, Y., Albu, M., Chen, X., Taipale, J., Hughes, T.R., and Weirauch, M.T. (2018). The Human Transcription Factors. Cell 175, 598–599.

53. Houlard, M., Romero-Portillo, F., Germani, A., Depaux, A., Regnier-Ricard, F., Gisselbrecht, S., and Varin-Blank, N. (2005). Characterization of VIK-1: a new Vav-interacting Kruppel-like protein. Oncogene 24, 28–38.

54. Shao, Z., Li, C., Wu, Q., Zhang, X., Dai, Y., Li, S., Liu, X., Zheng, X., Zhang, J., and Fan, H. (2022). ZNF655 accelerates progression of pancreatic cancer by promoting the binding of E2F1 and CDK1. Oncogenesis 11, 44.

55. Vaquerizas, J.M., Kummerfeld, S.K., Teichmann, S.A., and Luscombe, N.M. (2009). A census of human transcription factors: function, expression and evolution. Nat. Rev. Genet. 10, 252–263.

56. Oksuz, O., Henninger, J.E., Warneford-Thomson, R., Zheng, M.M., Erb, H., Vancura, A., Overholt, K.J., Hawken, S.W., Banani, S.F., Lauman, R., et al. (2023). Transcription factors interact with RNA to regulate genes. Mol. Cell 83, 2449–2463.e13.

57. He, C., Sidoli, S., Warneford-Thomson, R., Tatomer, D.C., Wilusz, J.E., Garcia, B.A., and Bonasio, R. (2016). High-Resolution Mapping of RNA-Binding Regions in the Nuclear Proteome of Embryonic Stem Cells. Mol. Cell 64, 416–430.

58. Yang, C., Zheng, J., Liu, X., Xue, Y., He, Q., Dong, Y., Wang, D., Li, Z., Liu, L., Ma, J., et al. (2020). Role of ANKHD1/LINC00346/ZNF655 Feedback Loop in Regulating the Glioma Angiogenesis via Staufen1-Mediated mRNA Decay. Mol. Ther. Nucleic Acids 20, 866–878.

59. Teng, Z., Yao, J., Zhu, L., Zhao, L., and Chen, G. (2021). ZNF655 is involved in development and progression of non-small-cell lung cancer. Life Sci. 280, 119727.

60. Jolma, A., Yan, J., Whitington, T., Toivonen, J., Nitta, K.R., Rastas, P., Morgunova, E., Enge, M., Taipale, M., Wei, G., et al. (2013). DNA-binding specificities of human transcription factors. Cell 152, 327–339.

61. Hernandez, N., and Weiner, A.M. (1986). Formation of the 3’ end of U1 snRNA requires compatible snRNA promoter elements. Cell 47, 249–258.

62. Latos, P.A., Goncalves, A., Oxley, D., Mohammed, H., Turro, E., and Hemberger, M. (2015). Fgf and Esrrb integrate epigenetic and transcriptional networks that regulate self-renewal of trophoblast stem cells. Nat. Commun. 6, 7776.

63. Yue, J., Lai, F., Beckedorff, F., Zhang, A., Pastori, C., and Shiekhattar, R. (2017). Integrator orchestrates RAS/ERK1/2 signaling transcriptional programs. Genes Dev. 31, 1809–1820.

64. Weissmann, F., Petzold, G., VanderLinden, R., Huis In‘tVeld, P.J., Brown, N.G., Lampert, F., Westermann, S., Stark, H., Schulman, B.A., and Peters, J.-M. (2016). biGBac enables rapid gene assembly for the expression of large multisubunit protein complexes. Proc. Natl. Acad. Sci. U. S. A. 113, E2564–9.

65. Bieniossek, C., Imasaki, T., Takagi, Y., and Berger, I. (2012). MultiBac: expanding the research toolbox for multiprotein complexes. Trends Biochem. Sci. 37, 49–57.

66. Evans, R., O’Neill, M., Pritzel, A., Antropova, N., Senior, A., Green, T., Žídek, A., Bates, R., Blackwell, S., Yim, J., et al. (2021). Protein complex prediction with AlphaFold-Multimer. bioRxiv, 2021.10.04.463034. 10.1101/2021.10.04.463034.

67. Mastronarde, D.N. (2003). SerialEM: A Program for Automated Tilt Series Acquisition on Tecnai Microscopes Using Prediction of Specimen Position. Microsc. Microanal. 9, 1182–1183.

68. Weis, F., and Hagen, W.J.H. (2020). Combining high throughput and high quality for cryo-electron microscopy data collection. Acta Crystallogr D Struct Biol 76, 724–728.

69. Punjani, A., Rubinstein, J.L., Fleet, D.J., and Brubaker, M.A. (2017). cryoSPARC: algorithms for rapid unsupervised cryo-EM structure determination. Nat. Methods 14, 290–296.

70. Bepler, T., Morin, A., Rapp, M., Brasch, J., Shapiro, L., Noble, A.J., and Berger, B. (2019). TOPAZ: A Positive-Unlabeled Convolutional Neural Network CryoEM Particle Picker that can Pick Any Size and Shape Particle. Microsc. Microanal. 25, 986–987.

71. Scheres, S.H.W., and Chen, S. (2012). Prevention of overfitting in cryo-EM structure determination. Nat. Methods 9, 853–854.

72. Goddard, T.D., Huang, C.C., Meng, E.C., Pettersen, E.F., Couch, G.S., Morris, J.H., and Ferrin, T.E. (2018). UCSF ChimeraX: Meeting modern challenges in visualization and analysis. Protein Sci. 27, 14–25.

73. Casañal, A., Lohkamp, B., and Emsley, P. (2020). Current developments in Coot for macromolecular model building of Electron Cryo-microscopy and Crystallographic Data. Protein Sci.

74. Sanchez-Garcia, R., Gomez-Blanco, J., Cuervo, A., Carazo, J.M., Sorzano, C.O.S., and Vargas, J. (2021). DeepEMhancer: a deep learning solution for cryo-EM volume post-processing. Commun Biol 4, 874.

75. Afonine, P.V., Poon, B.K., Read, R.J., Sobolev, O.V., Terwilliger, T.C., Urzhumtsev, A., and Adams, P.D. (2018). Real-space refinement in PHENIX for cryo-EM and crystallography. Acta Crystallogr D Struct Biol 74, 531–544.

76. Uhlén, M., Fagerberg, L., Hallström, B.M., Lindskog, C., Oksvold, P., Mardinoglu, A., Sivertsson, Å., Kampf, C., Sjöstedt, E., Asplund, A., et al. (2015). Proteomics. Tissue-based map of the human proteome. Science 347, 1260419.

